## Supplementary material for "The Utilization of Advance Telemetry to Investigate Important Physiological Parameters Including Electroencephalography in Cynomolgus Macaques Following Aerosol Challenge with Eastern Equine Encephalitis Virus": Supp. Figs. 1-4, Tables 1-2

### Supp. Figure 1

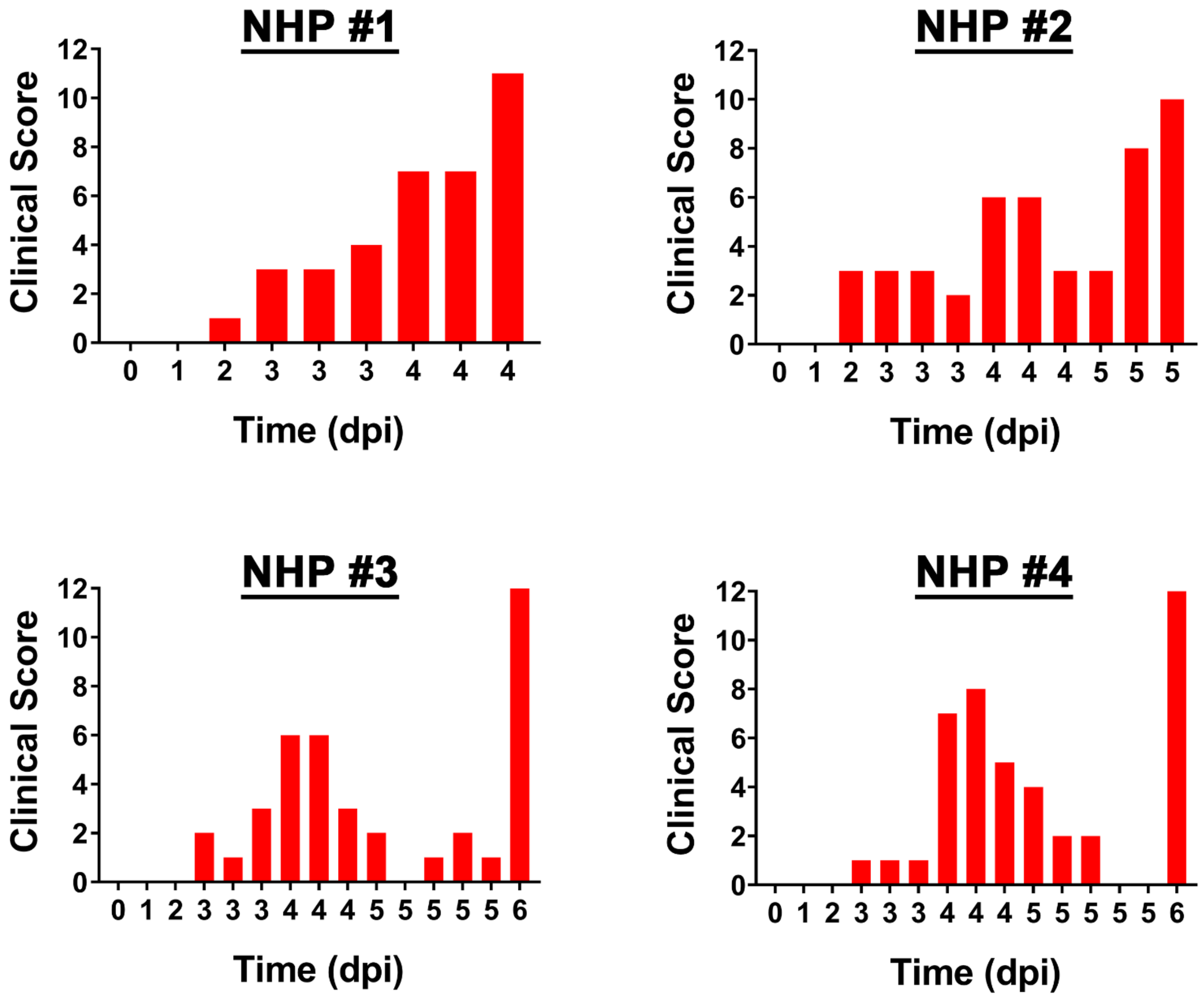

### Supp. Figure 2

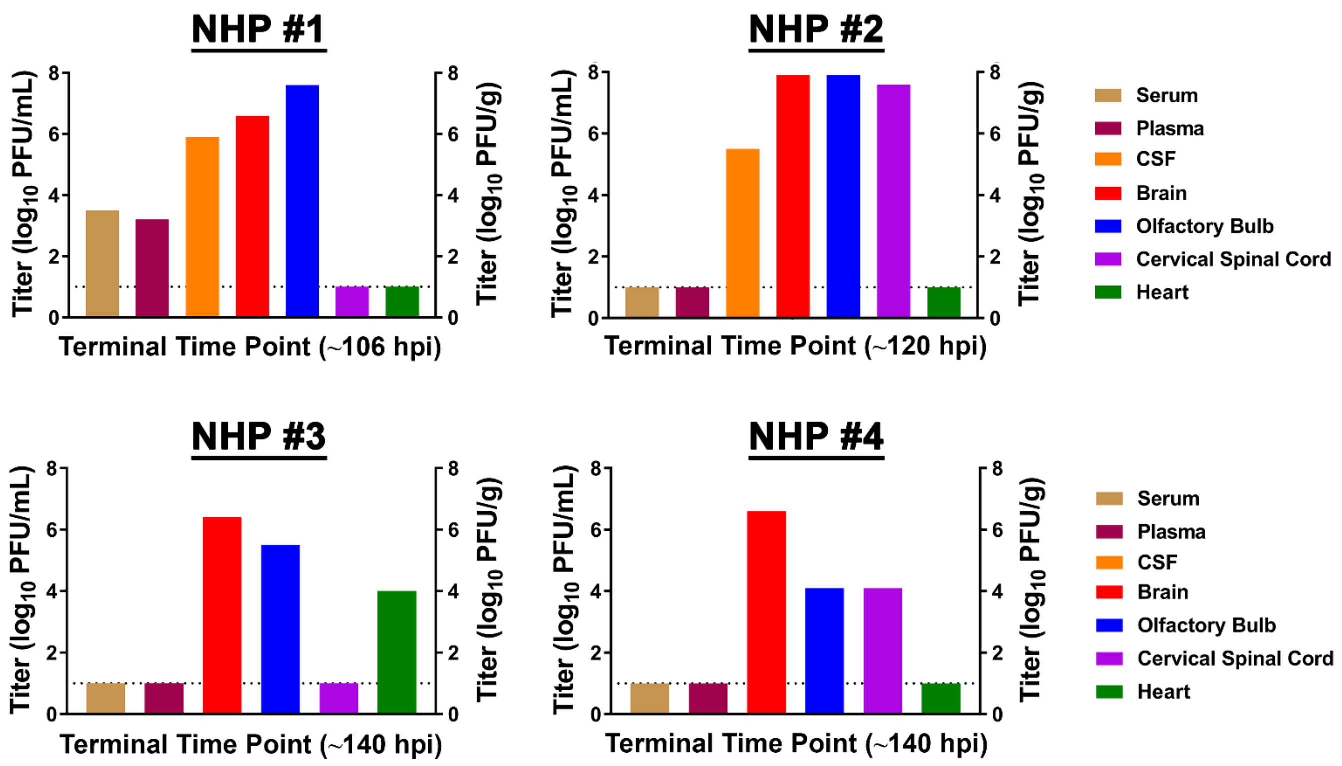

### Supp. Figure 3

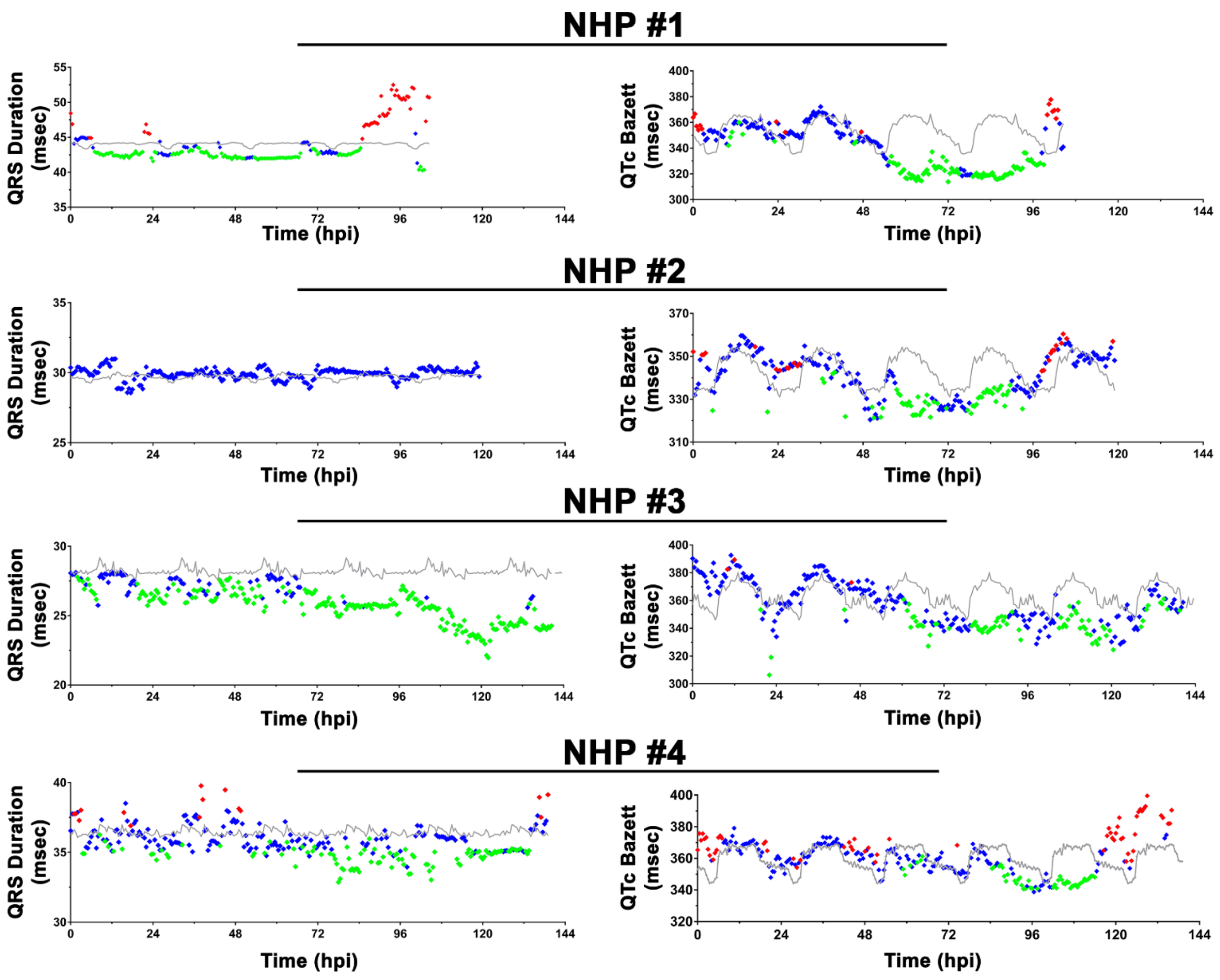

### Supp. Figure 4

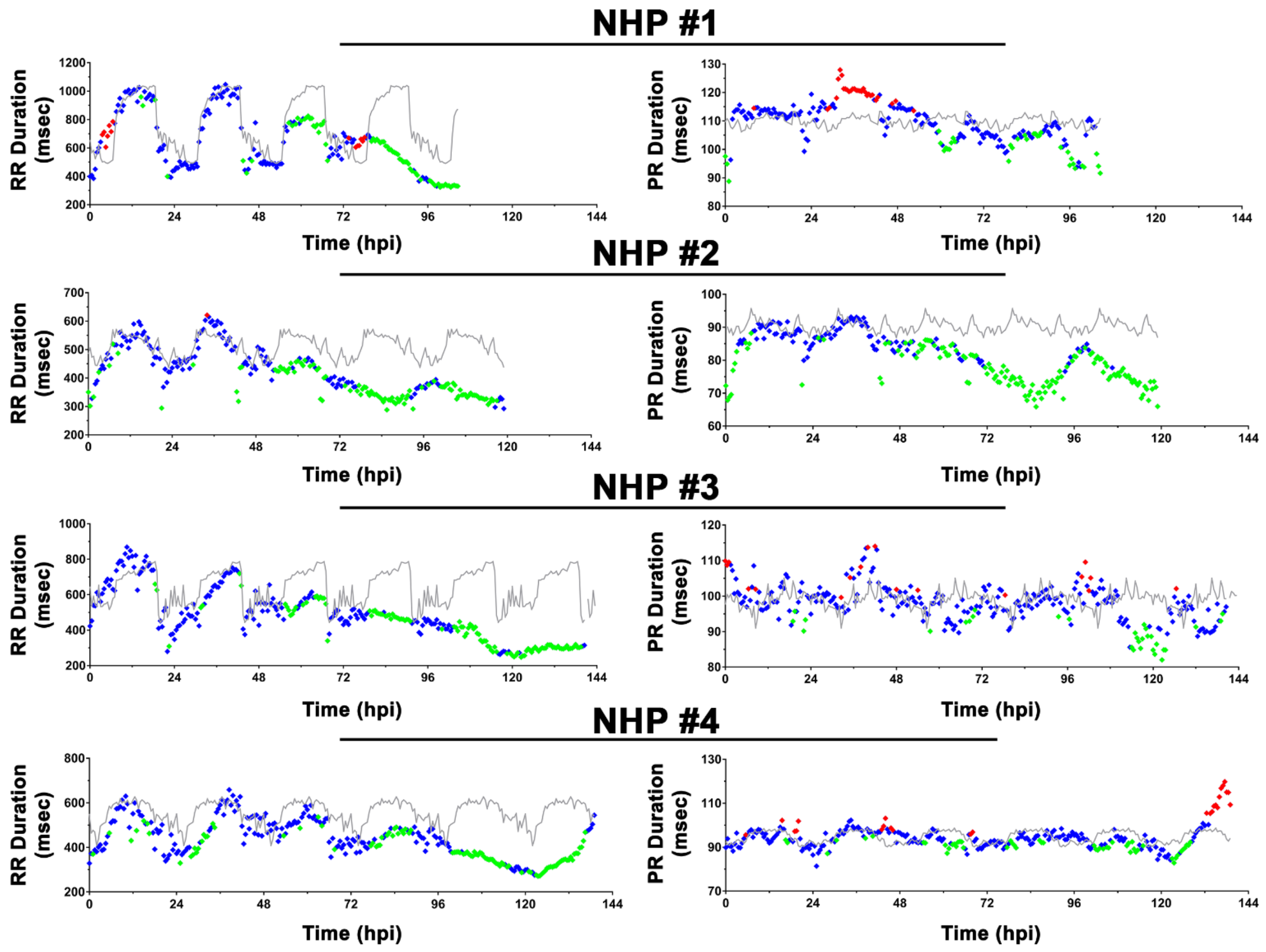

### Supp. Table 1

| Time (hpi) | NHP #1 | Time (hpi) | NHP #2 | Time (hpi) | NHP #3 | NHP #4 |
|---|---|---|---|---|---|---|
| 0-7 (Day) | Similar to baseline | 0-6 (Day) | Similar to baseline | 0-6 (Day) | Similar to baseline | Similar to baseline |
| 8-19 (Night) | Similar to baseline | 6-18 (Night) | Similar to baseline | 6-18 (Night) | Similar to baseline | ↓ Sleep,<br>↑ Activity |
| 20-31 (Day) | Similar to baseline | 18-30 (Day) | Similar to baseline | 18-30 (Day) | Similar to baseline | ↑ Sleep,<br>↓ Activity |
| 31-43 (Night) | Similar to baseline | 30-42 (Night) | Similar to baseline | 30-42 (Night) | Similar to baseline | ↓ Sleep,<br>↑ Activity |
| 43-55 (Day) | ↑ Restlessness,<br>↓ Fluid Consumption | 42-54 (Day) | Similar to baseline | 42-54 (Day) | ↓ Activity | ↑ Sleep,<br>↓ Activity |
| 55-67 (Night) | ↓↓ Sleep,<br>↑ Activity | 54-66 (Night) | ↓ Sleep,<br>↑ Activity | 54-66 (Night) | ↓ Sleep,<br>↑ Activity,<br>↓↓ Fluid Consumption | ↓ Sleep,<br>↑ Activity |
| 67-79 (Day) | ↓ Food and Fluid Consumption,<br>↓↓ Activity | 66-78 (Day) | ↓ Activity | 66-78 (Day) | ↑ Sleep,<br>↓ Activity,<br>↓↓ Fluid Consumption | ↓ Fluid Consumption,<br>↑ Sleep,<br>↓ Activity |
| 79-91 (Night) | ↓↓ Sleep,<br>↑ Activity | 78-90 (Night) | ↓↓ Sleep,<br>↑↑ Activity | 78-90 (Night) | ↓↓ Sleep,<br>↑ Activity,<br>↓↓ Fluid Consumption | ↓↓ Sleep,<br>↑ Activity |
| 91-103 (Day) | ↓↓↓Food and Fluid Consumption,<br>↓↓ Activity,<br>↑ Sialorrhea | 90-102 (Day) | ↓Food and Fluid Consumption,<br>↓↓ Activity | 90-102 (Day) | ↑ Sleep,<br>↓ Activity,<br>↓↓ Fluid Consumption | ↓↓ Fluid Consumption,<br>↑ Sleep Light,<br>↓ Activity |
| 106 (Night) | ↓↓↓Food and Fluid Consumption,<br>↓↓↓ Sleep,<br>↑ Sialorrhea,<br>***Onset of overt seizures***,<br>***Euthanasia*** | 102-114 (Night) | ↓↓↓Food and Fluid<br>↓↓Consumption,<br>↓↓ Activity,<br>↓↓↓ Sleep,<br>↑ Sialorrhea | 102-114 (Night) | ↓↓↓ Sleep,<br>↑ Activity,<br>↓↓↓ Fluid Consumption | ↓↓↓ Sleep,<br>↑ Activity |
| | | 120 (Day) | ↓↓↓Food and Fluid<br>↓↓Consumption,<br>↓↓ Activity,<br>↑ Sialorrhea,<br>***Onset of overt seizures***,<br>***Euthanasia*** | 114-126 (Day) | ↑ Sleep,<br>↓↓↓ Activity,<br>↓↓↓ Fluid Consumption | ↑ Sleep,<br>↓↓ Activity |
| | | | | 126-138 (Night) | ↓↓↓ Sleep,<br>↓↓↓ Activity,<br>↓↓↓ Fluid Consumption,<br>↑↑ Weakness/fatigue,<br>***Euthanasia*** ***(140 hpi)*** | ↓↓↓ Sleep,<br>↑↑↑ Activity followed by<br>↓↓↓ Activity,<br>↑ Weakness/fatigue,<br>***Onset of overt seizures***,<br>***Euthanasia*** ***(140 hpi)*** |

### Supp. Table 2

| Animal # | Onset of Fever (hpi) | Fever Duration (hrs) | Fever Hours | $\Delta T_{max}$ (°C) | Peak Temperature (°C) |
|---|---|---|---|---|---|
| NHP #1 | 52 | 50 | 160.8 | 4.2 | 41 |
| NHP #2 | 51 | 53.5 | 161.1 | 4 | 40.9 |
| NHP #3 | 56 | 45 | 127.3 | 3 | 40.1 |
| NHP #4 | 55 | 51 | 176.2 | 3.8 | 40.8 |
